# No evidence that proteome composition is associated with realised thermal limit and dietary niche breadth in butterflies

**DOI:** 10.1101/2024.12.04.626867

**Authors:** Fernanda S. Caron, Zuzanna Pietras, Arkan Eddine-Lomas, Rebecca von Hellfeld, Juliano Morimoto

## Abstract

Amino acids are the building blocks of proteins that perform essential physiological functions. Theory suggests that the proteome composition, the amino acid frequencies across all proteins in a genome, is associated with an organism’s optimal growth temperature, offering insights into species’ temperature limits. However, this hypothesis, based on prokaryotes, has not been tested in complex multicellular eukaryotes where many amino acids are strictly acquired through diet. Here, we analysed amino acid frequencies in the proteomes of orthologous and non-orthologous genes from 35 butterfly species to test for correlations with maximum observed temperatures and diet breadth. Using a comparative approach, we found no evidence that proteome composition correlates with temperature or diet breadth. Our findings suggest that animal proteome composition is likely shaped more strongly by energetic and biophysical constraints rather than by ecological factors.

## Introduction

The Anthropocene is marked by rising temperatures (Summerhayes and Zalasiewicz, 2018), climate extremes (Pradhan *et al*., 2022), and biodiversity loss (Dirzo *et al*., 2014; Johnson *et al*., 2017; Turvey and Crees, 2019). To survive, species must rapidly adapt to unpredictable conditions at the genomic, organismal, and behavioural levels, which is an unprecedented task in evolution (Bradshaw and Holzapfel, 2006; Gienapp *et al*., 2008; Hoffmann and Sgrò, 2011). Research has focused on understanding the correlations and causes associated with species’ response to, and interaction with, their environment, seeking biological patterns that can be used to conserve what is left of our biodiversity (Mawdsley, O’Malley and Ojima, 2009; Hannah, 2010). However, there are still many aspects of organismal biology and its relationship to the environment that remain unexplored (Merilä, 2012; Bozinovic and Pörtner, 2015).

One such case is the environmental effects on the proteome – i.e., the collection and frequency of amino acids from coding sequences of a genome (also referred to as ‘exome’) (Du *et. al*., 2018). Theory predicts that changes in amino acid frequencies across proteomes are associated with optimal growth temperatures of the species, and past empirical work has shown that mesophilic and thermophilic prokaryotes indeed have distinct amino acid frequencies in their proteomes (Dufton, 1997; Seligmann, 2003; Singer and Hickey, 2003; Tekaia and Yeramian, 2006; Swire, 2007). Work from our group has shown that amino acid frequencies in the proteomes of three domains of life (archaea, bacteria, and eukaryotes) respond to growth temperature, although this effect appears to be relatively small (Morimoto and Pietras, 2024). Nevertheless, these findings suggest that proteomes can be sensitive to temperature. To date, however, we lack studies to investigate the effects of temperature on proteome composition in multicellular eukaryotes (Roberts, 1999; but see e.g. Jensen *et al*., 2021).

Recent studies which focused on the proteome of multicellular eukaryotes have shown that the amino acid frequencies in fruit flies (*Drosophila melanogaster*) and mice (*Mus musculus*) contain information about dietary their requirements (Piper *et al*., 2017; Gómez Ortega *et al*., 2023). Matching the amino acid availability of the diet to the amino acid frequencies of the proteome improved reproduction without associated lifespan costs to in flies and improved growth rate in mice (Piper *et al*., 2017). This effect in flies was observed so long as the amino acid availability in the diet matched the absolute amino acid frequencies of the proteome, not the amino acid frequencies weighted by gene transcription (Gómez Ortega *et al*., 2023).

Feeding is one process among many through which organisms interact with their environments, and diet availability and quality is known to be affected by climate change (Stephens and Krebs, 1986; Simpson and Raubenheimer, 2012; Rosenblatt and Schmitz, 2016; Macdiarmid and Whybrow, 2019; European Food Safety Authority (EFSA) *et al*., 2020). Yet, feeding allows multicellular organisms to acquire essential amino acids which compose proteins and ultimately, the proteome (Simpson and Raubenheimer, 2012; Piper *et al*., 2017). In insect pollinators, it is known that temperature modulates availability of nutrients (Shrestha *et al*., 2016). Therefore, it is plausible, although yet untested, that proteomes with certain compositions belong to generalist (or specialist) species which can tolerate higher thermal limits. This would result in an association between proteome composition, diet breadth, and the maximum temperature in which species are encountered. Finding such patterns would be groundbreaking because proteome data is relatively cheap to collect and there are well-curated databases available for investigations, as opposed to time-consuming, labour-intensive experiments to ascertain diet breadth and thermal limits.

Here, we set out to test this and analysed the proteomes of 35 butterfly species obtained from the NCBI database. We tested whether amino acid frequencies in the proteome were associated with species’ diet breadth, the recorded maximum temperature where each species was observed (henceforth ‘climatic high temperatures’), or an interaction between them. Herbivorous insects such as butterflies are ideal models to test the effects of temperature and foraging on proteomes for three reasons. First, butterflies and other arthropods are particularly vulnerable to climate change (Harvey *et al*., 2023), have been declining at incredible pace (Habel *et al*., 2019; Warren *et al*., 2021), but are often overlooked in conservation policies (Duffus and Morimoto, 2022). Second, the effects of climate change and urbanisation on biodiversity loss appears to disproportionately affect diet specialist species, which may be more vulnerable to rapid and unpredictable changes in food supply or quality (Merckx and Van Dyck, 2019; Hulshof *et al*., 2024). And third, there are multiple fully annotated genomes available for butterflies and other insects, making the study of amino acid profiles across proteomes feasible at larger scales (Espeland *et al*., 2018; Liu *et al*., 2020; Kawahara *et al*., 2023). We first analysed whether the overall amino acid frequencies in the proteomes correlated with maximum temperature that species occur and species’ diet breadth and predicted that labile amino acids such as cysteine would have reduced frequencies for proteomes of species that are found in higher maximum temperatures (Seligmann, 2003; Singer and Hickey, 2003; Tekaia and Yeramian, 2006; Swire, 2007,

Morimoto and Pietras, 2024). Next, we separated the amino acid frequencies of orthologous and non-orthologous genes. Our rationale was that, due to differences in evolutionary patterns and selective pressures, these two classes of genes could give us different insights into the relationship between proteomes and species’ ecology. We predicted that because orthologous genes are often under selection to maintain ancestral functions across multiple species (Gabaldón and Kooni, 2013), they could be less likely to diversify to the extent that allows us to capture associations with species’ temperature limits. Conversely, non-orthologous genes could evolve and gain functions in specific lineages, providing a better subset of the proteome to identify correlations with species’ temperature limits. Our findings suggest that proteomes are imprinted by intrinsic (physiological needs) as opposed to extrinsic (ecological) responses.

## Materials and Methods

### Proteome analysis

The NCBI database from which the proteome for the 35 butterfly species were retrieved was accessed on or before January 6^th^, 2024 (Morimoto and Pietra, 2024). We retrieved proteome information for species which had complete and annotated reference sequence (RefSeq) genomes with suitable taxonomic identification in the NCBI database. This approach ensured that our estimates of amino acid frequencies were true representative of amino acids from coding sequences. The proteome fasta files were downloaded from FTP servers and were processed in the statistical software R version 4.3.2 (R Core Team, 2013) to estimate amino acid profiles. Because of the positive correlation between the number of redundant codons and the frequency of amino (Estimate: 0.0102, std error: 0.00032; t-value: 32.08; p < 0.001), we standardised our amino acid profiles, dividing amino acid frequency by the number of redundant codons. For all analysis, we normalised amino acid frequencies using the ‘scale()’ function in R with default parameters.

### Orthologous analysis

Inferring ortholog and non-ortholog genes from the proteomes of the studied species requires effective algorithms which form accurate gene trees. OrthoFinder is a recently developed orthologue inference algorithm that has innovated in providing more accurate ortholog inferences without sacrificing time costs (Emms and Kelly 2019). OrthoFinder’s algorithm was run on the collective lepidoptera proteomes and inferred orthologs from orthogroup trees it produced. Within OrthoFinder’s default algorithm, the sequence search method used was DIAMOND (Buchfink, Xie, and Huson 2015) and the orthogroup tree inference method used was DendroBlast (Kelly and Maini 2013). This method has been proven to achieve accurate results with low run time costs (Emms and Kelly 2015; Altenhoff et al. 2016). Amino acid frequency for each orthologous and non-orthologous gene in each species as well as for each species whole proteome was calculated for the phylogenetic comparative analysis.

### Phylogenetic reconstruction

To assess the correlation between Lepidoptera’s traits, we needed to reconstruct the phylogeny of the species in the present study. To do this, we obtained COI sequences from each species from GenBank (last accessed 20 August 2024; Table S1). Additionally, we obtained sequences for two outgroup species that allowed us to root the estimated phylogeny: *Cheumatopsyche brevilineata* and *Hydromanicus wulaianus*. The sequences were aligned using auto strategy in MAFFT online service v.7 (Kuraku et al., 2013; Katoh et al., 2019). Then, we reconstructed the phylogenetic relationships of the species on IQ-TREE v.2.3.6 (Minh et al., 2020). Simultaneously, we ran ModelFinder using the option -m MFP+MERGE to find the best evolutionary model for each codon position (Kalyaanamoorthy et al., 2017). We also ran an ultrafast bootstrap (Hoang et al., 2018) with 1000 replicates with branch lengths to assess phylogenetic uncertainty in the subsequent analyses. The resulting maximum likelihood tree did not show unusually long branches that could indicate poor alignment or obvious identification errors in some species. Finally, all bootstrap replicates were calibrated with PATHd8 (Britton et al., 2007). The calibrations used were obtained from Kawahara et al. (2023). As these authors used several distinct calibration schemes, we chose the calibration scheme that they used in their subsequent analyses while filtering (subsetting) the nodes we had present in our phylogeny (Table S2; Fig 1). We assigned a maximum age and a minimum age to each calibration, except for the oldest calibration. This is due to the requirement of PATHd8 that at least one calibration be fixed.

**Figure 1.**
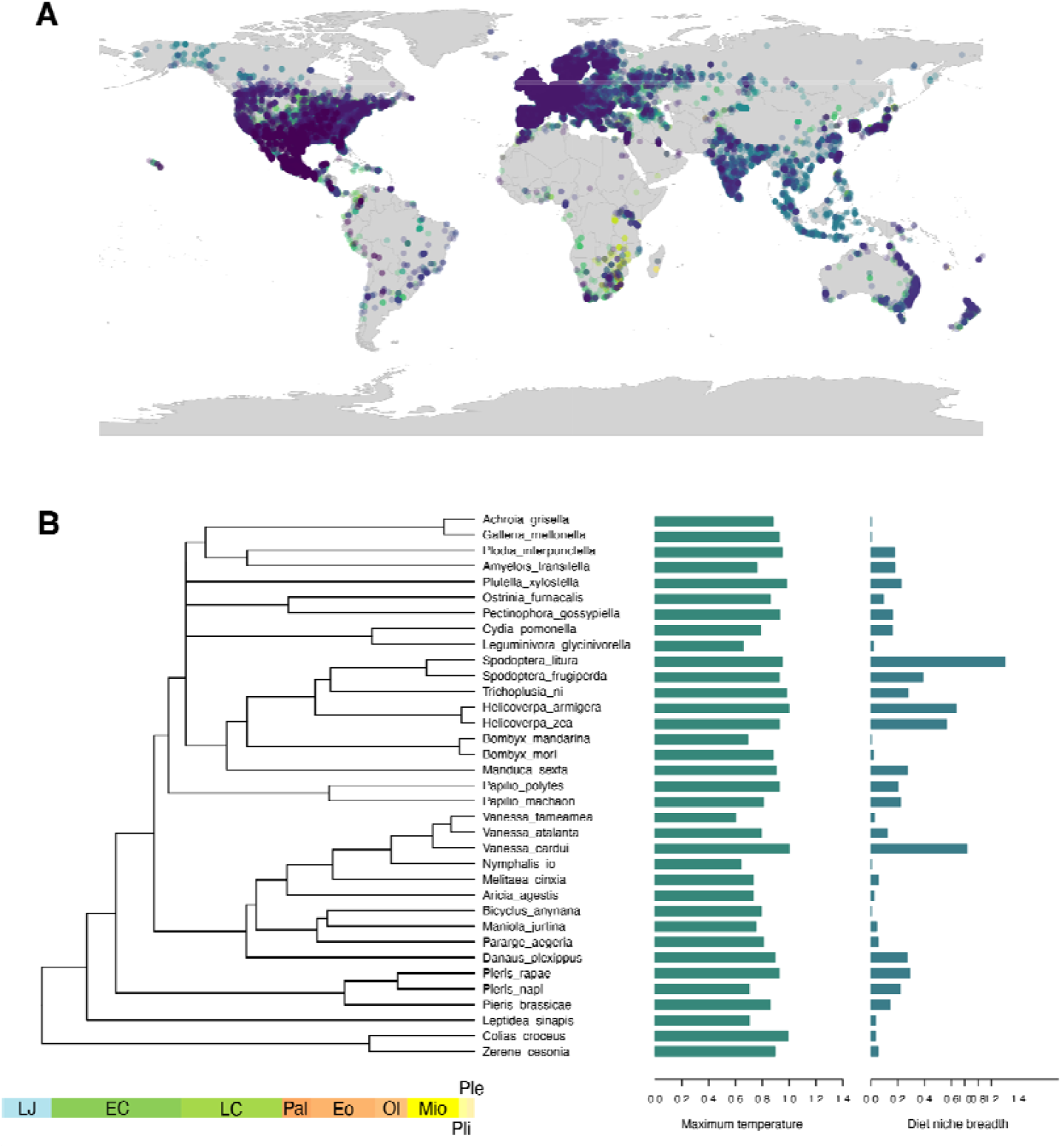
Ecological traits and phylogenetic relationships. (a) Coverage of our observation points for the 35 species from which maximum temperature and diet breadth were used to test associations with proteome composition. (b) Phylogenetic reconstruction of the relationship between the 35 species in our database (including 2 outgroups) along with the distribution of maximum temperature and dietary niche breadth of each species (right-hand side bars).

### Ecological and occurrence data

We retrieved and consolidated diet breadth data from two databases on March 6^th^ 2024: (1) HOSTS, a database on host plant for insects by the Natural History Museum in London (Robinson *et al*., 2010) and (2) DBIF, also a resource for recorded host plants for insects which have been recorded in Great Britain (Smith and Roy, 2008; Ward *et al*., 2019). There are limitations of using these public databases, such as the potential for taxonomic misidentification of either plants or insects, but they provide an invaluable resource to explore biological phenomena at larger scales (see e.g., Padovani *et al*., 2020). We estimated diet breadth at the genus level, that is, the number of different plant genus that a butterfly species has been recorded as using as a food source. Occurrences of each species were retrieved from GBIF (https://doi.org/10.15468/dl.q74xtz, 11th March 2024; Fig. 1). Climatic variables where each species occurs were obtained using the worldclim_global function with a resolution of 2.5 in the *geodata* package v.0.6-2 (Hijmans et al., 2023). For the subsequent analyses, we calculated the maximum temperature that each species has been found as a proxy of their thermal limit using the bio5 variable (maximum temperature of warmest month). Our rationale was that the maximum upper temperature where the species has been observed represented an upper limit on their thermal tolerance.

### Phylogenetic comparative analyses

We performed a Phylogenetic Least Squares Regression (PGLS) to assess the effect of diet and temperature on amino acid counts. Diet breadth and maximum temperature were log-transformed prior to the analyses. A PGLS analysis was done for the thermal limit and diet breadth, repeated for the 1000 topology replicates to account for phylogenetic uncertainty. We ran a PGLS for each amino acid separately as their frequencies vary widely. We repeated the PGLS three times considering the amino acid frequency of all genes pooled together at first, then only ortholog genes or non-ortholog genes. To control for multiple comparisons, we adjusted the p-value reported in this study by using the ‘p.adjust()’ function specifying the Benjamini & Hochberg (1995) correction method.

## Results

Our PGLS showed no evidence that individual amino acid frequencies in the proteome correlated with maximum temperature or diet breadth (Table 1). This was consistent for the analysis of the entire proteome (all genes), orthologous, and non-orthologous genes (Table 1). Together, these results show that proteome composition is not correlated with ecological traits related to thermal limit and diet breadth in the cosmopolitan butterfly species analysed in this study.

**Table 1.**
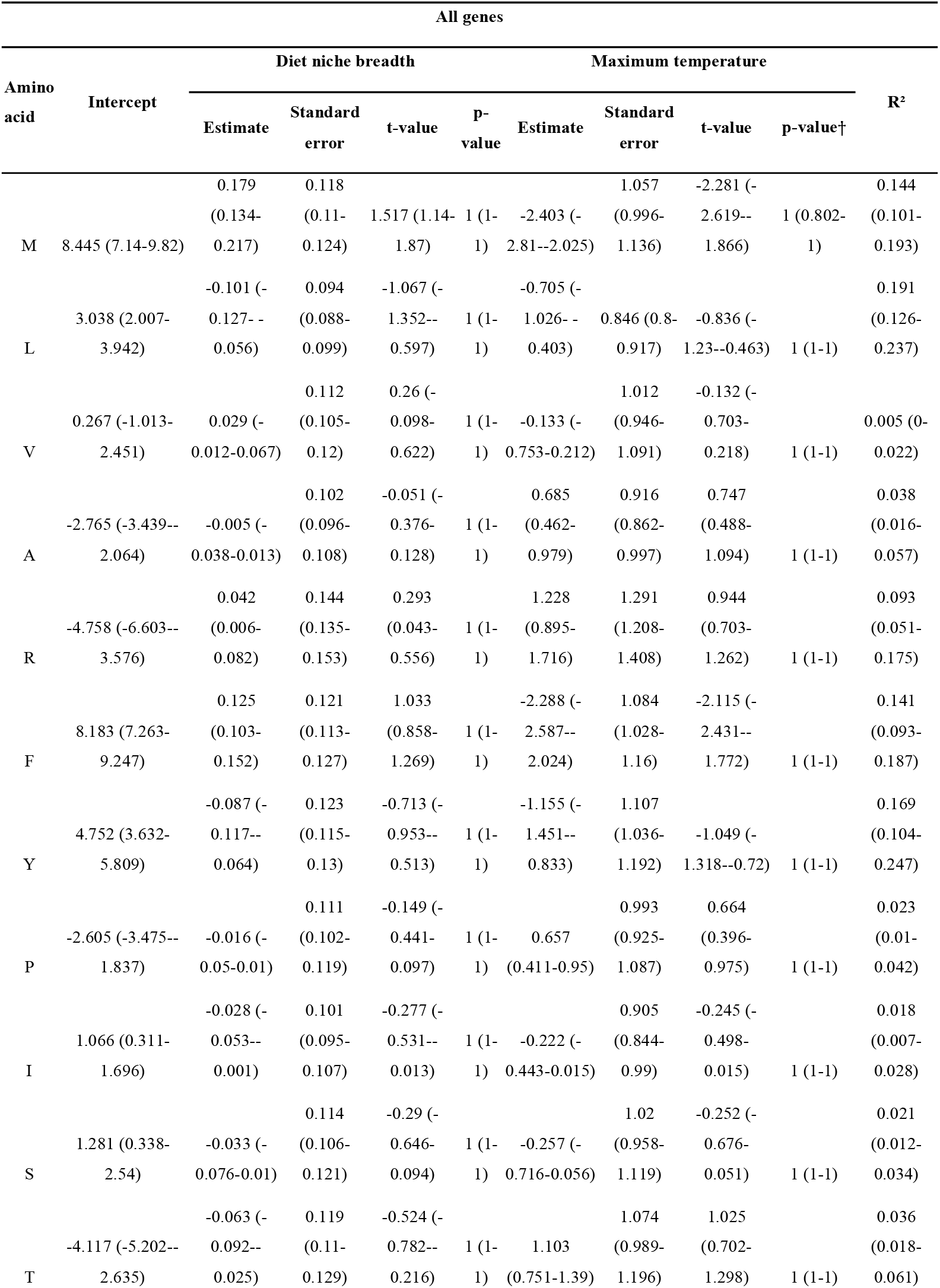
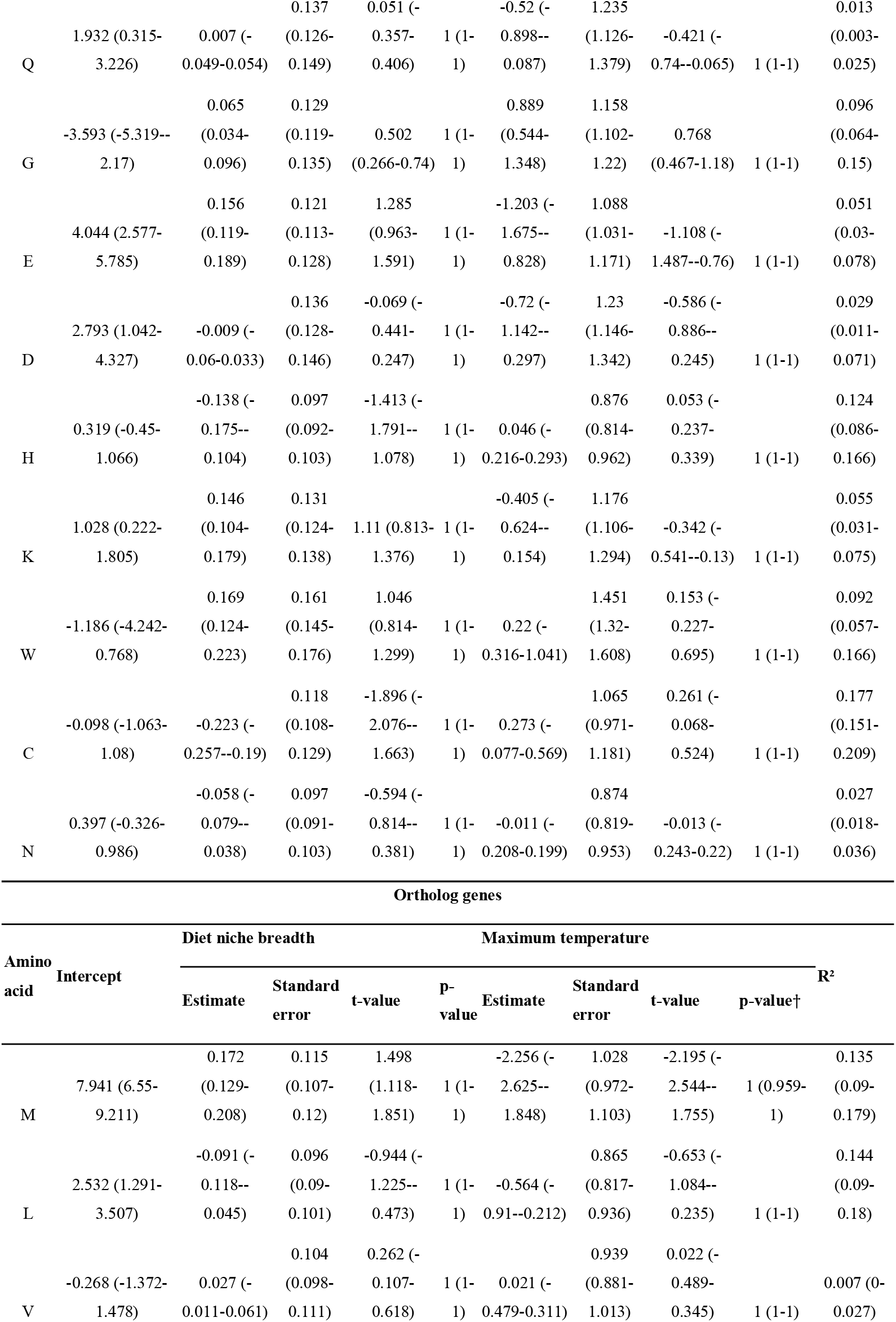
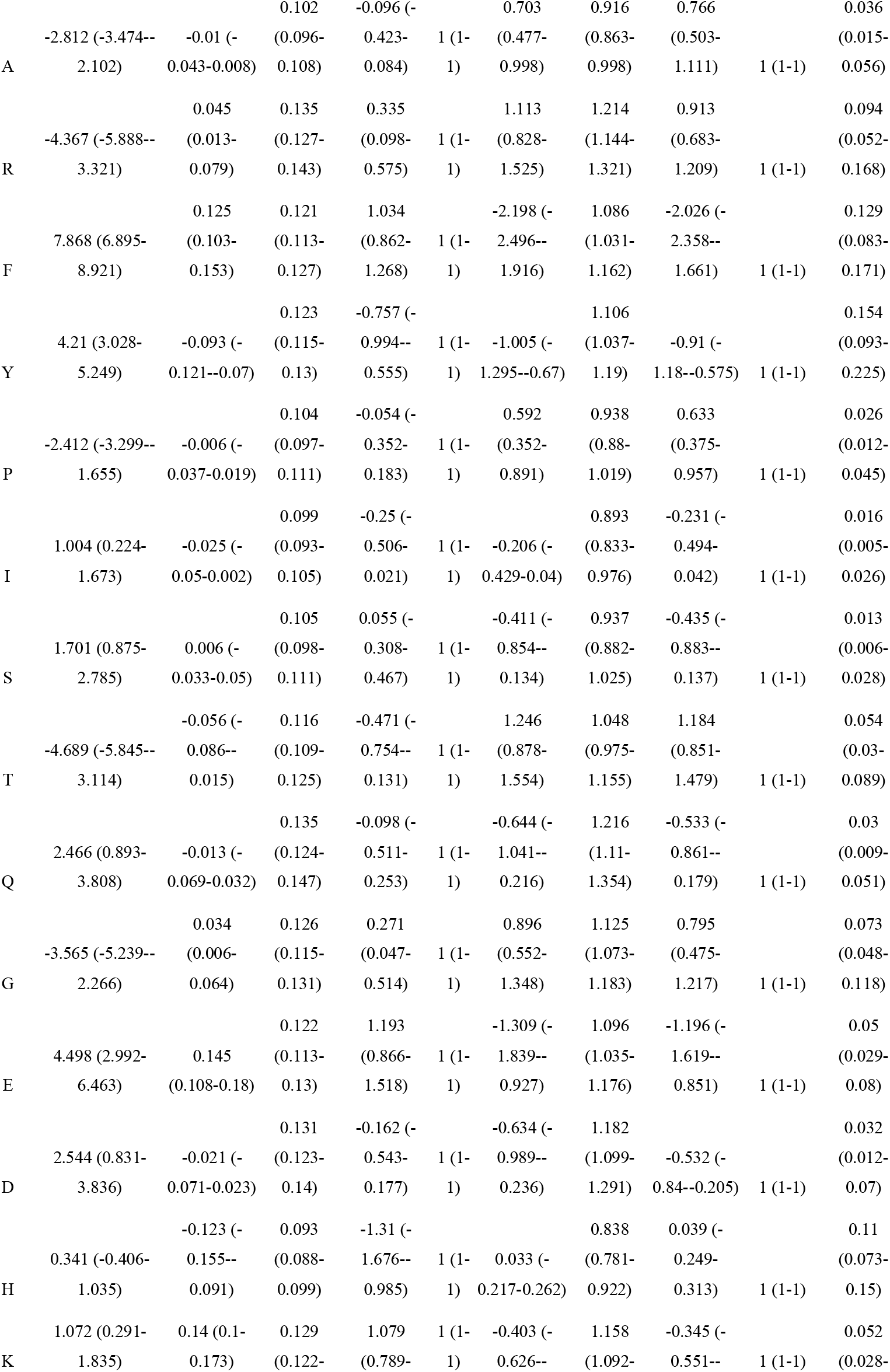
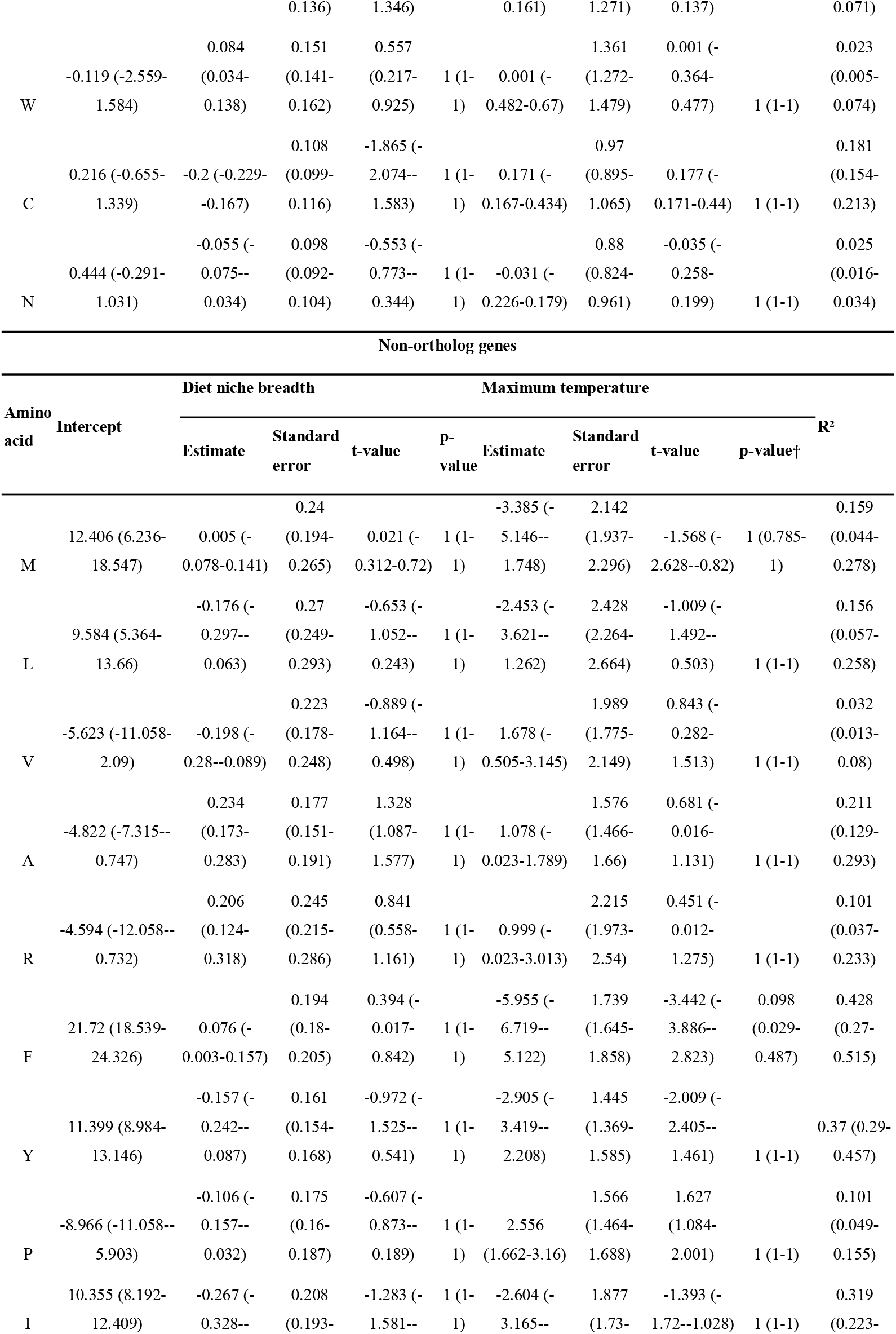
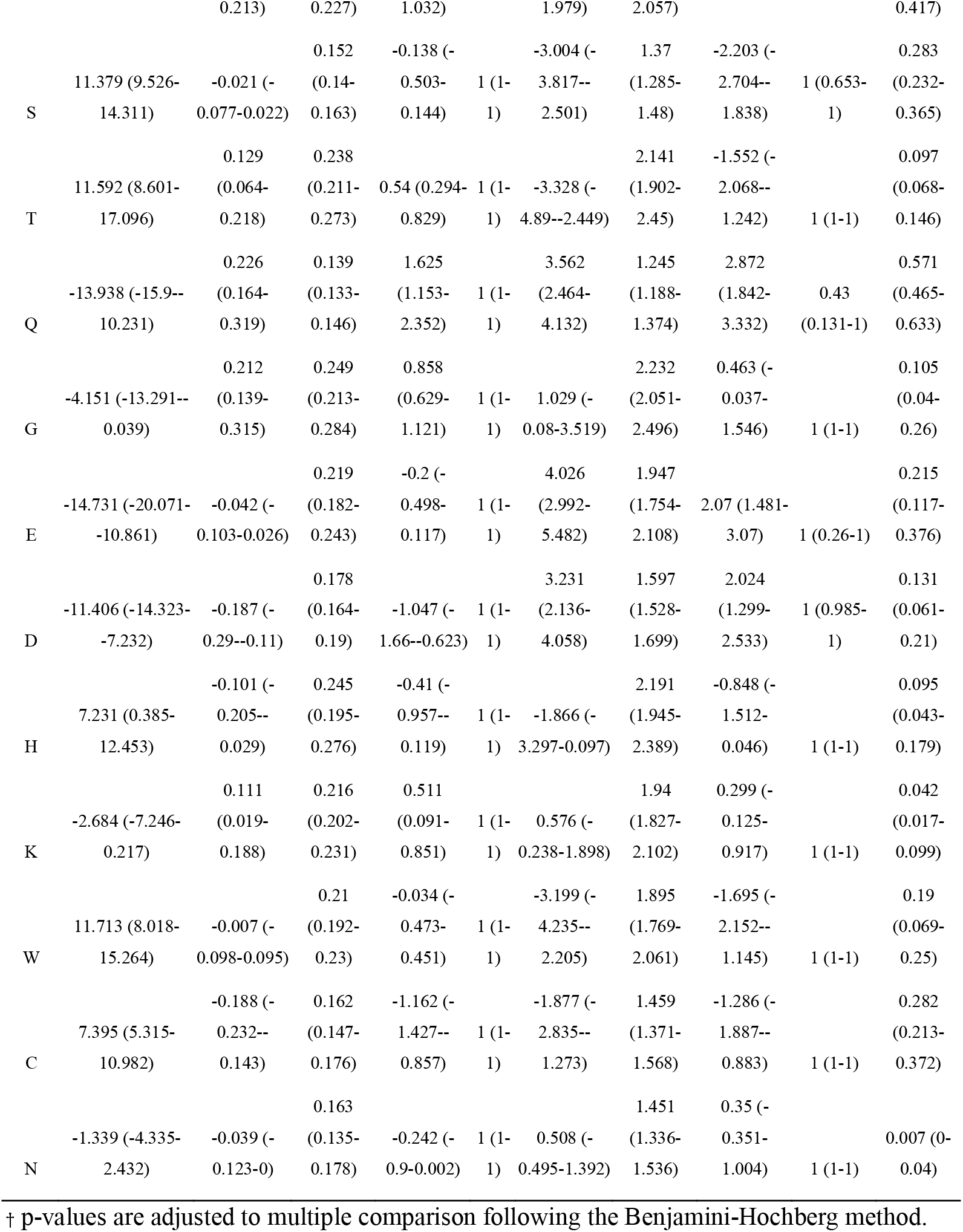
No evidence that proteome composition is associated with ecological traits in 35 butterfly species. Estimates obtained from PGLS. Values in parenthesis correspond to the interval of each parameter considering the topology of the trees (see Methods).

## Discussion

We present the first comprehensive test for the hypothesis that amino acid frequencies in the proteome are associated with temperature limits of complex multicellular organisms.

Specifically, we used a comparative approach to test whether the amino acid frequencies in the proteome of 35 butterfly species correlated with the maximum temperature that species are observed as well as with their dietary niche breadth. This hypothesis is derived from previous literature in prokaryotes which showed that the amino acid frequencies in the proteome are dependent upon the optimal growth temperature of a species, raising the intriguing possibility that the proteome contains information about the thermal ecology of the organism (Dufton, 1997; Chen & Nielsen, 2022; Seligmann, 2003; Singer and Hickey, 2003; Tekaia and Yeramian, 2006; Swire, 2007). Our most recent comprehensive study tested this hypothesis across three domains of life (archaea, bacteria, eukaryotes) and found supporting evidence that the frequency of thermolabile amino acids such as cysteine in the proteome are negatively associated with optimal growth temperature (Morimoto and Pietras, 2024). However, our past study focused on growth temperature and did not correlate growth temperature with species’ thermal limits. With the growing impacts of climate change, uncovering new relationships between species’ biology and ecology – such as the link between proteome, temperature, and diet studied here – is paramount to help understand species’ potential to tolerate increasing temperatures. There was no evidence that amino acid frequencies in the proteome were associated with maximum temperature or diet breadth of the 35 butterfly species. These results suggest that complex animal proteomes are likely shaped by energetic and biophysical constraints rather than by ecological factors portending to the temperature and diet niches which species can explore.

Amino acid frequencies in the proteome contain information about dietary needs underpinning life-history traits, but our data shows that this does not necessarily apply to broader ecological traits. For instance, Piper et al. (2017) showed that *D. melanogaster* and *M. musculus* fed diets with amino acids which were proportional to the amino acid frequencies of the proteome (“exome-matched” diets) optimises growth and reproduction without associated costs to lifespan. Gómez-Ortega et al. (2023) corroborated these in *D. melanogaster* and showed that diets which matched amino acid proteome composition were better at improving fecundity from both sexes compared with diets with amino acids with transcriptome-weighted proteome compositions, even though gene expression between sexes were substantial. This suggests that differential gene expression does not influence the broader information on amino acid needs of the organism and that the proteome inherently contains information about organismal nutritional needs above and beyond gene expression levels. Together, these findings show that proteome composition informs dietary needs for life history traits. Given this, we hypothesised that the proteome could also contain information about how organisms interact with their environment, under the assumption that the relationships between proteome composition, diet, and optimal growth temperature extended to other ecological traits. Here, we tested these relationships using analogous ecological traits to those found to correlate with proteome composition, namely maximum temperature (analogous to growth temperature) and dietary niche breadth (analogous to dietary needs). Yet, our results found no evidence that proteome composition was associated with either of these ecological traits. Thus, it appears that proteome compositions are linked to intrinsic (physiological) organismal needs (optimal growth temperature and diet to realise life-histories) but not with extrinsic (ecological) organismal responses (maximum temperature, dietary breadth). Whether this is a broader pattern across all multicellular organisms and across all ecological traits remains to be ascertained. Nonetheless, our findings show for the first time that, despite being linked to optimal growth temperature and dietary needs, proteome composition is not a good indicator of species thermal limits or dietary niches.

## Conclusion

Proteomes is a fundamental feature of biological systems and inform about the global amino acid requirements of a genome. Our work in butterflies suggest that proteomes are largely independent of observable ecological traits related to thermal limits and diet breadth. This opens the possibility that proteomes are primarily shaped by evolutionary constrains and does not contain signatures of ecological conditions in which species exist. This knowledge expands our understanding of evolutionary vs ecological constrains in proteome composition and partly contradicts evidence from prokaryotes and unicellular eukaryotes which showed that some ecological traits such as growth temperature influence proteome composition. Multicellular organisms are complex and might rely on more stable proteome compositions to achieve the multitude of physiological control and homeostasis across environments. Future studies in other taxonomic groups will provide valuable test of this hypothesis.

