## Supplementary File for "No evidence that proteome composition is associated with realised thermal limit and dietary niche breadth in butterflies"

### Authors’ Affiliations:

### ^1^ Programa de Pós-graduação em Ecologia e Conservação, Universidade Federal do Paraná, Curitiba, 82590-300, Brazil

### ^2^ Department of Physics, Chemistry and Biology (IFM), Linköping University, Sweden

### ^3^ School of Biological Sciences, University of Aberdeen, Cruickshank Building, Aberdeen, UK AB24 3UL

### ^4^ Institute of Mathematics, School of Natural and Computing Sciences, University of Aberdeen, Fraser Noble Building, Aberdeen, UK AB24 3UE

### Corresponding author information:

### Juliano Morimoto, PhD

### Institute of Mathematics, University of Aberdeen, AB24 3UE, Aberdeen

# +44 (0)1224 273218

###

#### **Table S1**. Accession numbers of the COI sequences used in the phylogenetic reconstruction.

| **Species** | **Accession numbers** |
| --- | --- |
| *Achroia grisella* | OM203125.1 |
| *Amyelois transitella* | KT692987.1 |
| *Aricia agestis* | AY350456.1 |
| *Bicyclus anynana* | KR139576.1 |
| *Bombyx mandarina* | AB070263.1 |
| *Bombyx mori* | PP993696.1 |
| *Colias croceus* | KM592967.1 |
| *Cydia pomonella* | JX407107.2 |
| *Danaus plexippus* | KC836923.1 |
| *Galleria mellonella* | KT750964.1 |
| *Helicoverpa armigera* | MZ397986.1 |
| *Helicoverpa zea* | OR722488.1 |
| *Leguminivora glycinivorella* | MZ506815.1 |
| *Leptidea sinapis* | DQ387045.1 |
| *Manduca sexta* | OP076895.1 |
| *Maniola jurtina* | AY090214.1 |
| *Melitaea cinxia* | NC_018029.1 |
| *Nymphalis io* | MZ322948.1 |
| *Ostrinia furnacalis* | ON645200.1 |
| *Papilio machaon* | NC_018047.1 |
| *Papilio polytes* | MZ188895.1 |
| *Pararge aegeria* | KJ547676.1 |
| *Pectinophora gossypiella* | ON479615.1 |
| *Pieris brassicae* | ON939547.1 |
| *Pieris napi* | OL332825.1 |
| *Pieris rapae* | OL790339.1 |
| *Plodia interpunctella* | KT428892.1 |
| *Plutella xylostella* | JF911819.1 |
| *Spodoptera frugiperda* | KU877172.1 |
| *Spodoptera litura* | JQ647918.1 |
| *Trichoplusia ni* | MK714850.1 |
| *Vanessa atalanta* | HQ734886.1 |
| *Vanessa cardui* | EF683677.1 |
| *Vanessa tameamea* | PP319022.1 |
| *Zerene cesonia* | KM046838.1 |
| *Cheumatopsyche brevilineat*a (ourgroup) | KX385010.1 |
| *Hydromanicus wulaianus* (outgroup) | KF717095.1 |

### Table S2. Calibration scheme used in the phylogenetic reconstruction based on Kawahara et al. (2023).

| **Calibration point** | **Minimum age** | **Maximum age** | **Median age** |
| --- | --- | --- | --- |
| Nymphalidae | - | - | 78.2 |
| Nymphalinae | 38.9 | 48.6 | - |
| Satyrinae | 54.0 | 58.7 | - |
| Papilioninae | 49.8 | 70.5 | - |
| Coliadinae | 36.0 | 44.1 | - |
| Pierinae | 44.5 | 49.9 | - |
